# Is Actin Filament and Microtubule Growth Reaction- or Diffusion-Limited?

**DOI:** 10.1101/406090

**Authors:** J. Pausch, G. Pruessner

## Abstract

Inside cells of living organisms, actin filaments and microtubules self-assemble and dissemble dynamically by incorporating actin or tubulin from the cell plasma or releasing it into their tips’ surroundings. Such reaction-diffusion systems can show diffusion- or reaction-limited behaviour. However, neither limit explains the experimental data: while the offset of the linear relation between growth speed and bulk tubulin density contradicts the diffusion limit, the surprisingly large variance of the growth speed rejects a pure reaction limit. In this Letter, we accommodate both limits and use a Doi-Peliti field-theory model to estimate how diffusive transport is perturbing the chemical reactions at the filament tip. Furthermore, a crossover bulk density is predicted at which the limiting process changes from chemical reactions to diffusive transport. In addition, we explain and estimate larger variances of the growth speed.

---

Microtubules and actin filaments are structures that polymerise by incorporating and releasing their diffusively moving building blocks, tubulin and actin. They are responsible for growth, shape, movement, and transport processes, among others, and can span the entire cell (1). Their dynamics have been studied intensively experimentally^1^ and theoretically.^2^ However, many questions about their dynamics remain debated or completely unanswered. This includes: Is their growth limited by diffusion to their tip (13,15) or by the chemical reaction rates for incorporation (12,15)? How can the large variance of their growth be explained (2)?

In experiments^1^, the filament growth speed ⟨ *v* ⟩ can be measured as a function of the bulk density *ζ* of tubulin or actin. An effective incorporation coefficient *k*_on_ and effective release rate *k*_off_ are determined as parameters of a linear fit of growth speed data over bulk concentration *ζ*

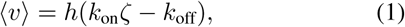

where *h* is the effective growth length per incorporated particle. There are two mean field approaches for modeling the filament growth speed ⟨ *v* ⟩.

The first approach assumes that diffusive transport is quicker than the chemical reactions(7,9-12) and that polymerisation is therefore *reaction-limited*. This effectively infinite diffusivity *D* implies that particle concentration is homogeneous, and in particular, that there is no significant depletion close to the filament tip. Thus, filament growth is determined by two Poisson processes, incorporation with rate *λζ* and release with rate *τ*, each of which is associated with a step of length *h*. The two competing processes create a Skellam distribution (16) with expected growth speed ⟨*v*⟩ _*R*_ and effective diffusion constant *D*_eff_

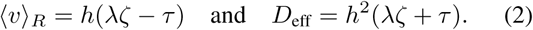

^1^See Table S1 in the Supplemental Information of (2) for microtubule assembly rates. For actin filaments, assembly rates can be found in (3–6).

^2^Theoretical work includes (7–13) for microtubules and (4, 8, 14) for actin filaments.

However, in comparison with experiments (2, 4, 5), *D*_eff_ is too small. Furthermore, it implies an independence of the growth speed from the viscosity of the medium, which was rejected experimentally for actin filaments (15) and microtubules (17). Thus, a purely reaction-limited behaviour is rejected.

The second approach assumes that transport by diffusion is slower than the chemical reactions (13,15) and that selfassembly is therefore *diffusion-limited*. This implies that, due to the incorporation into the tip, the building blocks are depleted locally. The growth speed is determined by the diffusive flux to the tip. If the protein concentration *c*(*x, t*) follows a steady state diffusion equation 0 = *D*Δ*c*(*x, t*), the particle flux *J* through the absorbent reaction sphere of radius *R* and the growth length *h* determine the growth speed ⟨*v* ⟩_*D*_, which is equivalent to Smoluchowski coagulation (18)

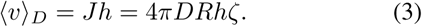

However, this limit cannot accommodate a release rate as any released particle would immediately be reabsorbed before diffusion can transport it away from the reaction surface.

Furthermore, using this approach with typical parameters of microtubule assembly, only a small reduction of the bulk tubulin density to 89% is found at the reaction surface (13). It therefore is not completely absorbent, as is theoretically suggested in (19). According to the Stokes-Einstein equation (20) for small Reynolds numbers, diffusion-limited growth implies that viscosity and incorporation rate are inversely proportional (21) without offset. Tested in (15) and (17), small offsets for the growth of microtubules and the barbed actin filament ends are found, while a significant offset is found for the pointed ends of actin filaments. Thus, filament growth cannot be perfectly diffusion-limited either. Furthermore, this model is not probabilistic and therefore, it is not clear how to derive a variance of the growth speed.

The diffusion(Eq. (3)) and reaction-limited (Eq. (2)), as well as the measured growth speed are schematically depicted in Fig. 1. Measured growth and shrinking speeds will always be slower than both limits. When shrinking, the chemically possible shrinking speed is not attained as diffusion slows down transport away from the tip, leading to a locally higher particle density. It can be read off the plot as the density at which the reaction-limited is equal to the measured speed (dotted line). Analogously, when growing, the reaction-limited speed is not attained either because diffusion fails to maintain the bulk particle density around the tip. The local density at the tip can be read off again as the density at which the reaction-limited speed is equal to the measured one (dotted line). In principle, there are three cases: the purely reaction limited case A, the mixed case B, where a crossover from reaction-limited to diffusion-limited behaviour occurs at the crossover bulk density *ζ*_*×*_, and case C (not shown), where shrinking does not occur and the growth is limited by diffusion at all particle densities.

**Figure 1:**
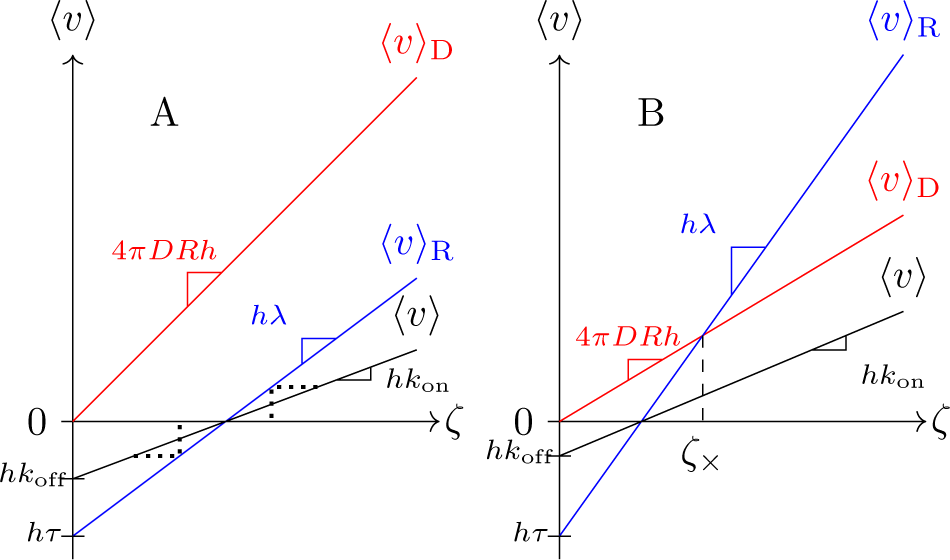
Filament growth speed ⟨*v* ⟩ as function of the bulk density *ζ* of its building blocks. The diffusion-limited ⟨*v* ⟩ _*D*_ (red), the reaction-limited ⟨*v* ⟩ _*R*_ (blue), and the real growth speed ⟨*v* ⟩ (black) are shown. A: The system is reactionlimited, B: Reaction-limited growth changes to diffusionlimited growth at the crossover bulk density *ζ*_*×*_. Local densities at the tip can be read off as the bulk density at which reaction-limited growth leads to the same speed (dotted line).

Theoretically, progress can be made by going beyond mean field theory which allows to calculate how chemical reactions are perturbed by diffusive transport of the reactants. Here, filament self-assembly is modelled on a threedimensional lattice and described by a master equation (Supplementary Material, Eq. (A4)). Following work by Doi (22) and Peliti (23), this model is transformed into a field theory. The derivation of the field theory is outlined briefly in the Supplementary Material, in Sections A2 and A3.

In our model, the building blocks and the filament tip are represented by fields, which are interpreted as time-dependent probability distributions of their positions. Due to the field representation, they do not have a size. However, the finite size of the proteins is an important element of the step-wise filament growth. Therefore, our model is set up on a hybrid, three dimensional space: While particles move in continuous space ℛ^3^, the filament tip is restricted to a discrete line 𝒵 with lattice constant *h*, overlaying the *z*-axis in ℛ^3^, see Fig. 2. The persistence length *P* and flexural rigidity *K* of the filament we model are thus effectively infinite. This is a good approximation for microtubules (*P ≈* 5200*µm*, *K ≈* 2 10^-^_23_*Nm*_2_), while for actin filaments, this approximation is slightly worse (*P ≈* 18*µm*, *K ≈* 7 10^-^_26_*Nm*_2_) (24). As we are modelling only a single polymer instead of the 13 microtubule protofilaments or the 2 actin filament strands, we interpret the lattice spacing *h* as the effective growth step length.

**Figure 2:**
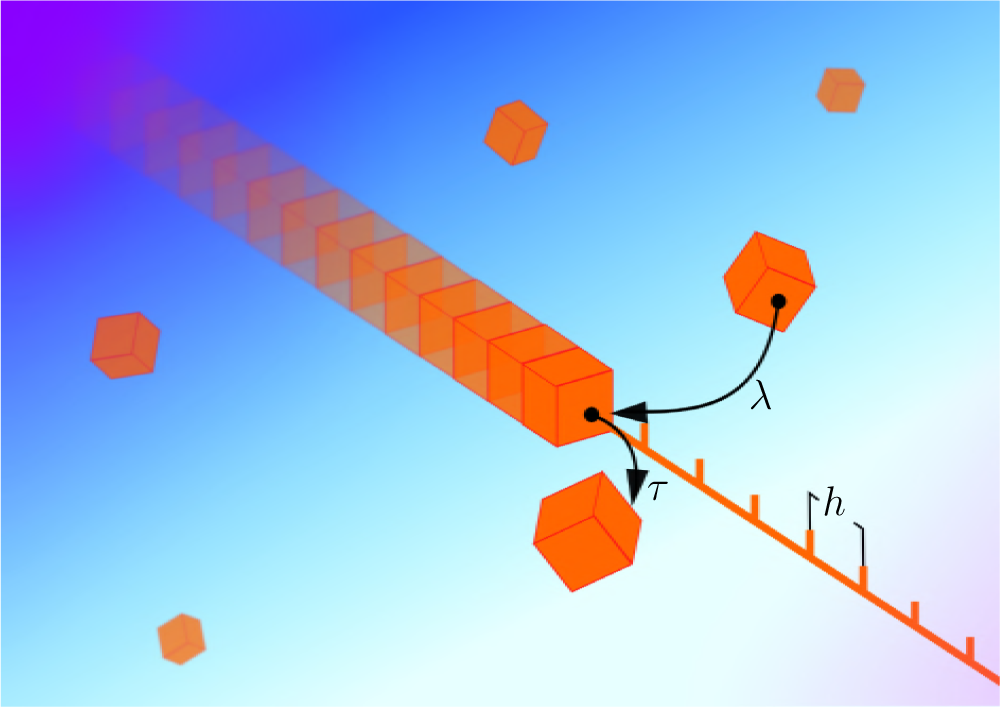
Schematic of the filament self-assembly process. Its building blocks move diffusively in ℛ^3^. The filament tip is on a lattice with spacing *h*. Particles can be incorporated in the filament (coefficient *λ*) and released by it (rate *τ*).

Each of the fields exists as an annihilation field and a creation field: *ϕ*(*x, t*) and *ϕ*^*†*^(*x, t*) for the building blocks, as well as *Ψ*_*j*_(*t*) and 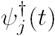 for the filament tip. The creation field initiates a single particle or filament tip at the specified position and time, whereas the annihilation field measures their number at the point stated. Creation fields often appear as Doi-shifted fields (22), e.g 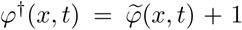. In addition, the particle annihilation field is shifted to measure deviations from the bulk density *ζ*, i.e 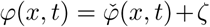. Between creation and annihilation, the system evolves by the stochastic processes included in the model.

There are six microscopic processes in our model. The units of the corresponding coefficients, denoted by [*…*], are written as monomials of *T* (time) and *L* (length).

- particle diffusion with constant *D*, [*D*] = *T*^-^_1_*L*_2_
- actin or tubulin absorption by the filament tip with coefficient *λ* and subsequent movement of the tip by distance *h* in the +*z* direction, [*λ*] = *T*^-^_1_*L*_3_
- particle release from the filament tip with rate *τ* and subsequent movement of the tip by distance *h* in the *-z* direction, [*τ*] = *T*^-^_1_
- actin/tubulin creation with coefficient *γ*, [*γ*] = *T*^-^_1_*L*^-^_3_
- extinction of actin/tubulin with rate *r*, [*r*] = *T*^-^_1_
- extinction of the filament tip with rate, *ϵ* [*ϵ*] = *T*^-^_1_

Creation and extinction of the building blocks is balanced such that a constant bulk density *ζ* = *γ/r* is created. The two extinction processes are included in the field theory to enforce causality. After calculations, we let parameters *γ, r* and tend to zero while keeping the ratio *γ/r* = *ζ* constant. Thus, the spontaneous extinction and creation are removed while a bulk density remains included.

The interaction part of *𝒜* has the form

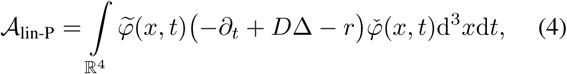

where the time dependency is omitted for better readability.

The different parts of the interaction describe the following processes:

(a) the filament grows, i.e. the filament tip moves one step in the positive *z* direction upon incorporating an actin/tubulin particle;

(b) as particles are incorporated into the tip, its density is reduced locally, resulting in anticorrelations of the tip and the particle density;

(c) particle density is reduced by incorporation into the tip;

All of the processes above are reflected in the action functional 𝒜of our model which splits up into a bilinear part and an interaction part 𝒜= 𝒜_lin_ + 𝒜_int_. The diffusion and extinction of particles is represented in the particle bilinear part

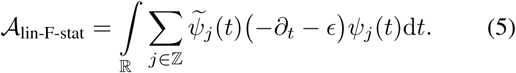

where Δ is the spatial Laplace operator.

The filament tip is stationary without the processes of incorporation or release of tubulin. It is described by

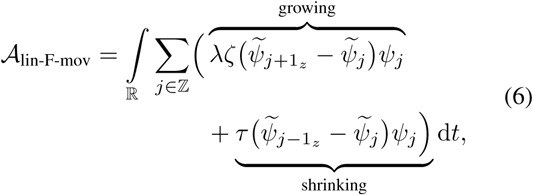

However, due to incorporation and release, the bilinear part includes jumps in steps of 1_*z*_

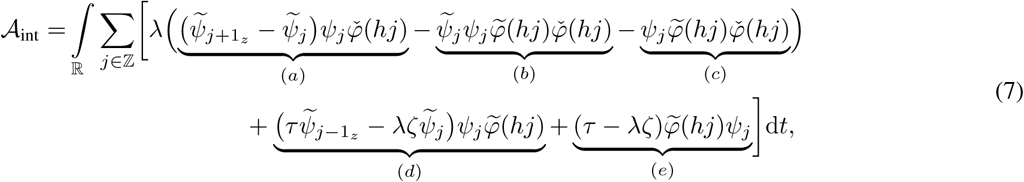

where we omitted the time dependence of the fields for better readability. The first part corresponds to filament growth, while the second part describes shrinking of the filament. Jumps on the lattice are indicated by *±*1_*z*_.

All three bilinear actions together make up *𝒜*_lin_ = *𝒜*_lin-P_+ *𝒜*_lin-F-stat_+ *𝒜*_lin-F-mov_.

(d) in the presence of a tip, the particle density is increased by spontaneous release (*τ*) and decreased by incorporation (*λζ*), leading to corresponding correlations of tip and particle densities;

(e) *τ*: particle density is increased because the filament releases a particle; *λζ*: particle density is decreased because the filament incorporates a particle.

Field-theoretic propagations and interactions are schematically represented by Feynman diagrams. Particle propagation is drawn as a straight red line, filament propagation is depicted as a curly blue line, and interactions are illustrated as vertices, see Fig. 3.

**Figure 3:**
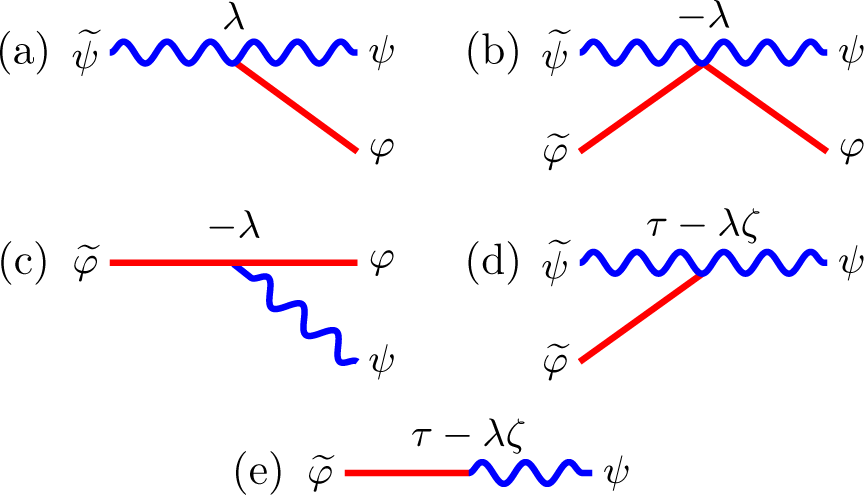
The stochastic processes that appear in 𝒜_int_, Eq. (7), are represented as amputated vertices in Feynman diagrams. Curly blue lines represent filaments, while straight red lines stand for their building blocks, actin or tubulin. All Feynman diagrams are read from right to left.

The interplay between propagation and interaction in a system governed by the action𝒜 can be calculated using the path integral. The system may be initialised by placing a filament tip at position *hj*_0_= 0 at time *t*_0_= 0. Then, the system evolves and particle concentrations, the filament tip positions, or moments of their distributions can be measured at a later point in time. In general, if the observable that we want to measure is represented by a combination of fields 𝒪 (*t*), then its time-dependent, spatial probability distribution is given by the following path integral (see e.g. (25) for a detailed review)

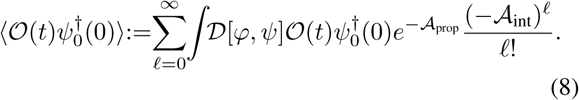

Formally, this integral is summing all variations of all fields involved of all stochastic processes possibly occurring. The path integral is normalised such that

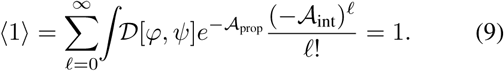

The *𝓁*-th term of the series is the contribution of all processes with *𝓁* interactions.^3^ The distribution of the filament’s tip position *j* is given by 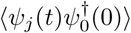. The expected growth speed and its variance are then determined by calculating the filament’s expected position and its variance after time *t* In the following, we consider three approximations of 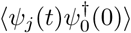: Firstly, 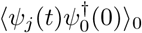 is the reaction-limited

^3^These interactions are in the field-theoretic sense. In fact, the term for *𝓁* = 0 already includes an arbitrary number of chemical interactions.

distribution, cutting the sum in Eq. (8) at *𝓁*= 0, which results in only one Feynman diagram shown in Eq. (10a). Secondly, 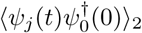 is terminating the sum at *𝓁*= 2, resulting in two diagrams shown in Eq. (10a) and (10b). Thirdly, 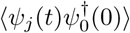 is the Dyson sum, which contains terms of all orders but selects only those Feynman diagrams whose loops are arranged daisy-chain-like, shown in Eq. (10):

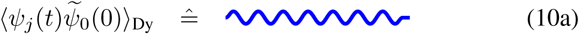

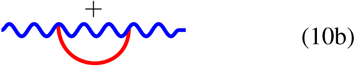

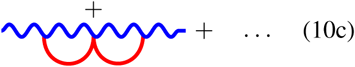

*A priori*, it is not clear which truncation of the path integral is a good approximation of the observable. However, a good agreement of the approximate result with experimental data indicates that the processes which were not included in the calculation rarely occur under experimental conditions.

The second and third approximation for the average filament growth speed are

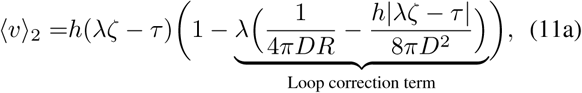

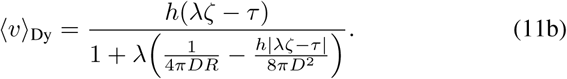

The approximation ⟨*v*⟩_0_is given by ⟨*v*⟩_2_without the loop correction term. The derivation of Eq. (11) is outlined in the Supplementary Material, in Section A5.

The loop correction term takes into account that, due to diffusive transport, the reaction-limited speed is reduced further because the locally depleted particles have to reach the reaction sphere of radius *R*. Considering the first term of the loop correction, if the diffusion is strong, the chemical reactions are less hindered by slow transport and the growth speed is closer to its reaction limit. This diffusion correction is itself corrected in the second term of the loop correction which describes how quickly the tip reaches regions that are less depleted. It predicts a non-linear dependence of the growth speed ⟨*v*⟩ on the bulk particle density *ζ*, which so far has not been observed experimentally. Therefore, we assume 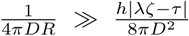 and ignore the latter in the following. Thus, given the diffusion-limited growth coefficient 4*πDR* and the observed *k*_on_ and *k*_off_, Eq. (1), we can calculate the reaction-limited *λ* as well as the offset *τ*

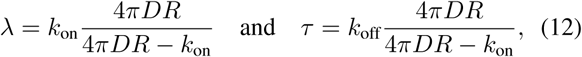

which is equivalent to the results in (26). It follows, that the observed growth coefficient *k*_on_ is smaller than the reactionand the diffusion-limited growth coefficients, i.e. 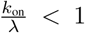 and 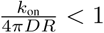, as shown in Fig. 1.

Furthermore, there is a bulk particle density *ζ*_*×*_ at which diffusion becomes the defining limitation in comparison to reaction (Fig. 1, B):

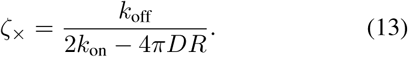

For *k*_off_ = 0, if 4*πDR >* 2*k*_on_, then growth is reaction-limited otherwise it is diffusion-limited.

One of the open questions for microtubule and actin filament growth is the origin of large fluctuations (2,4). In part, they can be explained as an overlap of the fluctuations of the reaction and diffusion processes. This is quantified by the effective diffusion constant (derived in SI, Sec. A6)

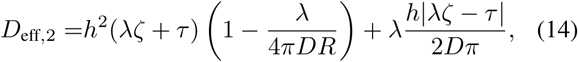

which is larger than the reaction-limited effective diffusion in Eq. (2).

In conclusion, the field theoretic model allows us to calculate how the reaction limit of microtubule and actin filament growth is undermined due to imperfect diffusive supply of tubulin or actin. It also predicts larger growth fluctuations compared to a model purely based on reactions. Given the overlap of fluctuations due to chemical reactions and diffusive transport, it is likely for the filament growth speed to exhibit correlations in time, the study of which would be compelling for future research.

The authors thank Robert Endres, Guillaume Salbreux and Thomas Surrey for very helpful discussions.

G.P. conceived the problem. G.P. and J.P. jointly developed the theory. J.P. carried out calculations and wrote the manuscript, guided by G.P.

## References

1. Boal, D., 2012. Mechanics of the Cell. Cambridge University Press.

2. Gardner, M. K., B. D. Charlebois, I. M. Jánosi, J. Howard, A. J. Hunt, and D. J. Odde, 2011. Rapid Microtubule Self-Assembly Kinetics. Cell 146:582–592.

3. Pollard, T. D., 1984. Polymerization of ADP-Actin. J. Cell Biol. 99:769–777.

4. Fujiwara, I., S. Takahashi, H. Tadakuma, T. Funatsu, and S. Ishiwata, 2002. Microscopic analysis of polymerization dynamics with individual actin filaments. Nat. Cell Biol. 4:666–673.

5. Kuhn, J. R., and T. D. Pollard, 2005. Real-Time Measurements of Actin Filament Polymerization by Total Internal Reflection Fluorescence Microscopy. Biophys. J. 88:1387–1402.

6. Fujiwara, I., D. Vavylonis, and T. D. Pollard, 2007. Polymerization kinetics of ADP- and ADP-P*i*-actin determined by fluorescence microscopy. Proc. Natl. Acad. Sci. Unit. States Am. 104:8827–8832.

7. Mitchison, T., and M. Kirschner, 1984. Dynamic Instability of Microtubule Growth. Nature 312:237–242.

8. Frieden, C., 1985. Actin and Tubulin Polymerization: The Use of Kinetic Methods to Determine Mechanism. Annu. Rev. Biophys. Bio. 14:189–210.

9. Carlier, M. F., R. Melki, D. Pantaloni, T. Hill, and Y. Chen, 1987. Synchronous oscillations in microtubule polymerization. Proc. Natl. Acad. Sci. Unit. States Am. 84:5257–5261.

10. der Chen, Y., and T. Hill, 1987. Theoretical studies on oscillations in microtubule polymerization. Proc. Natl. Acad. Sci. Unit. States Am. 84:8419–8423.

11. Walker, R., E. O’Brien, N. Pryer, M. Soboeiro, W. Voter, H. Erickson, and E. Slamon, 1988. Dynamic Instability of Individual Microtubules Analyzed by Video Microscopy: Rate Constants and Transition Frequencies. J. Cell Biol. 107:1437–1448.

12. Bayley, P., 1993. Why Microtubules Grow and Shrink. Nature 363:309.

13. Odde, D. J., 1997. Estimation of the Diffusion-Limited Rate of Microtubule Assembly. Biophys. J. 73:88–95.

14. Guo, K., J. Shillcock, and R. Lipowsky, 2009. Self-Assembly of actin monomers into long filaments: Brownian dynamics simulations. J. Chem. Phys. 131:1–11.

15. Drenkhahn, D., and T. D. Pollard, 1986. Elongation of Actin Filaments is a Diffusion-limited Reaction at the Barbed End and Is Accelerated by Inert Macromolecules. J. Biol. Chem. 261:12754–12758.

16. Skellam, J. G., 1946. The Frequency Distribution of the Difference Between Two Poisson Variates Belonging to Different Populations. J. Roy. Stat. Soc. 109:296.

17. Wieczorek, M., S. Chaaban, and G. J. Brouhard, 2013. Macro-molecular Crowding Pushes Catalyzed Microtubule Growth to Near the Theoretical Limit. Cell. Mol. Bioeng. 6:383–392.

18. v. Smoluchowski, M., 1917. Versuch einer mathematischen Theorie der Koagulationskinetik kolloider Lösungen. Z. Phys. Chem. 92:129–168.

19. Collins, F. C., 1950. Diffusion in chemical reaction processes and in the growth of colloid particles. J. Colloid Sci. 5:499–505.

20. Sutherland, W., 1905. A Dynamical Theory of Diffusion for Non-Electrolytes and the Molecular Mass of Albium. Phil. Mag. 9:781–785.

21. Berg, O. G., and P. H. von Hippel, 1985. Diffusion-controlled macromolecular interactions. Ann. Rev. Biophys. Chem. 14:131–160.

22. Doi, M., 1976. Second quantization representation for classical many-particle system. J. Phys. A: Math. Gen. 9:1465–1477.

23. Peliti, L., 1985. Path integral approach to birth-death processes on a lattice. J. Phys.-Paris 46:1469–1483.

24. Gittes, F., B. Mickey, J. Nettleton, and J. Howard, 1993. Flexural Rigidity of Microtubules and Actin Filaments Measured from Thermal Fluctuations in Shape. J. Cell Biol. 120:923–934.

25. Taüber, U. C., 2014. Critical Dynamics. Cambridge University Press.

26. Noyes, R. M., 1961. Effects of diffusion rates on chemical kinetics. Prog. React. Kinet. 1:129–160.

